# A comprehensive catalogue of regulatory variants in the cattle transcriptome

**DOI:** 10.1101/2020.12.01.406280

**Authors:** Shuli Liu, Yahui Gao, Oriol Canela-Xandri, Sheng Wang, Ying Yu, Wentao Cai, Bingjie Li, Ruidong Xiang, Amanda J. Chamberlain, Erola Pairo-Castineira, Kenton D’Mellow, Konrad Rawlik, Charley Xia, Yuelin Yao, Pau Navarro, Dominique Rocha, Xiujin Li, Ze Yan, Congjun Li, Benjamin D. Rosen, Curtis P. Van Tassell, Paul M. Vanraden, Shengli Zhang, Li Ma, John B. Cole, George E. Liu, Albert Tenesa, Lingzhao Fang

## Abstract

Characterization of genetic regulatory variants acting on the transcriptome of livestock is essential for interpreting the molecular mechanisms underlying traits of economic value and for increasing the rate of genetic gain through artificial selection. Here, we build a cattle Genotype-Tissue Expression atlas (cattle GTEx, http://cgtex.roslin.ed.ac.uk/) as part of the pilot phase of Farm animal GTEx (FarmGTEx) project for the research community based on publicly available 11,642 RNA-Seq datasets. We describe the landscape of the transcriptome across over 100 tissues and report hundreds of thousands of genetic associations with gene expression and alternative splicing for 24 major tissues. We evaluate the tissue-sharing patterns of these genetic regulatory effects, and functionally annotate them using multi-omics data. Finally, we link gene expression in different tissues to 43 economically important traits using both transcriptome-wide association study (TWAS) and colocalization analyses to decipher the molecular regulatory mechanisms underpinning such agronomic traits in cattle.

## Introduction

Genome-wide association studies (GWAS) have identified thousands of genetic variants associated with complex traits in human and livestock populations^1,2^. As the majority of these variants are non-coding, the characterization of the molecular mechanisms by which such variants affect complex traits has been extremely challenging. Indeed, in human genetics, it would have been impossible without projects such as the Genotype-Tissue Expression (GTEx) project that has characterized genetic effects on the human transcriptome and paved the way to understanding the molecular mechanisms of human variation^3^.

However, livestock genomic resources lag behind human genomic resources, and to date, no study has systematically explored the regulatory landscape of the transcriptome across a wide range of tissues. GWAS signals of agronomic traits are significantly enriched in regulatory regions of genes expressed in trait-relevant tissues in cattle^4-6^, but experiments to dissect genetic variation in gene expression have generally been small, both in terms of the number of individuals and tissues. For instance, a few of studies have explored the expression/splicing quantitative trait loci (e/sQTL) in blood^7^, milk cells^7^, muscle^8^ and mammary gland in cattle^9^. Here, we describe the largest and most comprehensive study of the regulatory landscape of any livestock species by analyzing 11,642 publicly available cattle RNA-Seq datasets, representing over 100 different tissues and cell types. We combined all of these data and make the results freely and easily accessible to the research community through a web portal (http://cgtex.roslin.ed.ac.uk/).

There has been a recent exponential growth in the number of RNA-Seq samples made publicly available in cattle (Fig. S1a), but these data have never been gathered in one collection and processed uniformly before. Here, we present a pipeline to uniformly integrate 11,642 public RNA-Seq datasets and identify eQTLs and sQTLs for 24 important cattle tissues with sufficient sample sizes (n > 40). The latter is facilitated by calling variants directly from the RNA-Seq reads and imputing to sequence level using a large multi-breed reference panel^10^, in a similar process to that used with human data^11^. Next, we conducted *in silico* analyses to annotate eQTLs and sQTLs with a variety of publicly available omics data in cattle, including DNA methylation, chromatin states, and chromatin conformation characteristics. Finally, we integrated gene expression with a large GWAS of 27,214 dairy bulls and 43 cattle traits *via* both transcriptome-wide association study (TWAS) and colocalization analyses to detect genes and variants associated with these economically important traits. The cattle Genotype-Tissue Expression (cattle GTEx) atlas will serve as a primary source of reference for cattle genomics, breeding, adaptive evolution, veterinary medicine, and comparative genomics.

## Results

### Data summary

We analyzed 11,642 public RNA-Seq datasets from 8,653 samples, yielding ∼200 billion clean reads (Table S1). Summary distributions of sequencing platform, read type (single/paired reads), clean read number, read length, sex, age, and mapping rate across samples show that the quality of these publicly available data is acceptable for the following analyses (Fig. S1b-h)^11^. We kept 7,180 samples with clean read > 500,000 and mappingv rate > 60% for subsequent analyses, representing 114 tissues from 46 breeds and breed combinations (Fig. S1i, Table S1). Holstein was the most represented breed (35.5% of all samples), reflecting its global economic value. A total of 1,831 samples (21%) had no breed records, but that information could be predicted from the genotypes called from RNA-Seq data. We grouped the 114 tissues into 13 categories based on known biology and the 46 breeds into six sub-species, with *Bos taurus* representing 87% of all samples (Table S1). To investigate the tissue-specificity of DNA methylation and to functionally annotate QTLs, we also analyzed 18 newly generated and 126 existing whole-genome bisulfite sequence (WGBS) samples from 21 cattle tissues, producing ∼73 billion clean reads with an average mapping rate of 71% (Table S2).

### General characteristics of transcriptome across samples

As expected, the number of expressed genes (Transcripts per Kilobase Million, TPM > 0.1) increased with the number of clean reads across samples. However, we observed a plateau at 50 million clean reads (Fig. S2a) where we only detected ∼60% of 27,607 Ensembl annotated genes. Only 61 genes were not expressed in any of the samples, and 33 of them (54.10%) were located in unplaced scaffolds, with significantly (*P* < 0.05) shorter gene length, fewer exons, higher CG density, and lower sequence constraints than expressed genes (Fig. S2b-f). Similarly, we detected more alternative splicing events with increasing numbers of clean reads across samples (Fig. S2g). However, we did not detect splicing events for 874 genes in any sample, which also exhibited significantly shorter gene length, fewer exons, lower expression, and lower sequence constraints than spliced genes (Fig. S2h-k). Furthermore, 27% of them were snRNAs, snoRNAs and rRNAs that play important roles in RNA splicing^12^ (Fig. S2l). Genes without splicing events were significantly enriched in the integral component of membrane and G-protein coupled receptor signaling pathways (Fig. S2m). We found that ∼25% of CpG sites in the entire genome were not covered at 5× in any of the WGBS samples, even if these had more than 300 million clean reads, partially due to bisulfite treatment and PCR amplification bias (Fig. S3a). These CpG sites were enriched in gene deserts (e.g., telomeres) with significantly higher CG density than the CpG sites captured by the WGBS (Fig. S3b-d).

We called a median of 21,623 SNPs from all RNA-Seq samples (Fig. S4a), and then imputed each sample up to 3,824,444 SNPs using a multi-breed reference population of 3,310 animals^10^. We validated the imputation accuracy by comparing SNPs derived from RNA-Seq with those called from whole-genome sequence (WGS) in the same individuals, including Holstein, Limousin and Angus, and the concordance rates were over 99% (Fig. S4b, c, and Table S3). We also compared the imputed genotypes from RNA-Seq data with those imputed using 50K SNP array genotypes in a subset of 109 Holstein animals. Although there was a depletion of high-quality (DR^2^ > 0.80) imputed intergenic variants amongst SNPs imputed from RNA-Seq data (Fig. S4d), the DR^2^ values of SNPs imputed from RNA-Seq were similar to those imputed from SNP-array along 1Mb up-/down-stream of gene body (covering genomic regions for *cis*-QTL mapping) (Fig. S4e). In addition, the correlation of genotype counts between imputed SNPs from RNA-Seq data and those from SNP array was around 0.80 (Fig. S4f). For the subsequent *cis*-QTL mapping, we focused on 24 tissues with greater than 40 individuals, and this encompassed 5,911 samples. After removing duplicated samples within each tissue (Fig. S4g), we retained 4,889 individuals.

We found that clusters of samples derived from both gene expression and alternative splicing in the uniformly analyzed data accurately recapitulated tissue types (Fig. 1a, b), reinforcing the quality and therefore their utility for our follow-up analysis. For instance, all the muscle samples from over 40 projects clustered together. Similar to expression and splicing, DNA methylation profiles also recapitulated tissue types (Fig. 1c). However, when clustering based on imputed genotypes, as expected, samples clustered by sub-species (Fig. 1d).

**Fig. 1.**
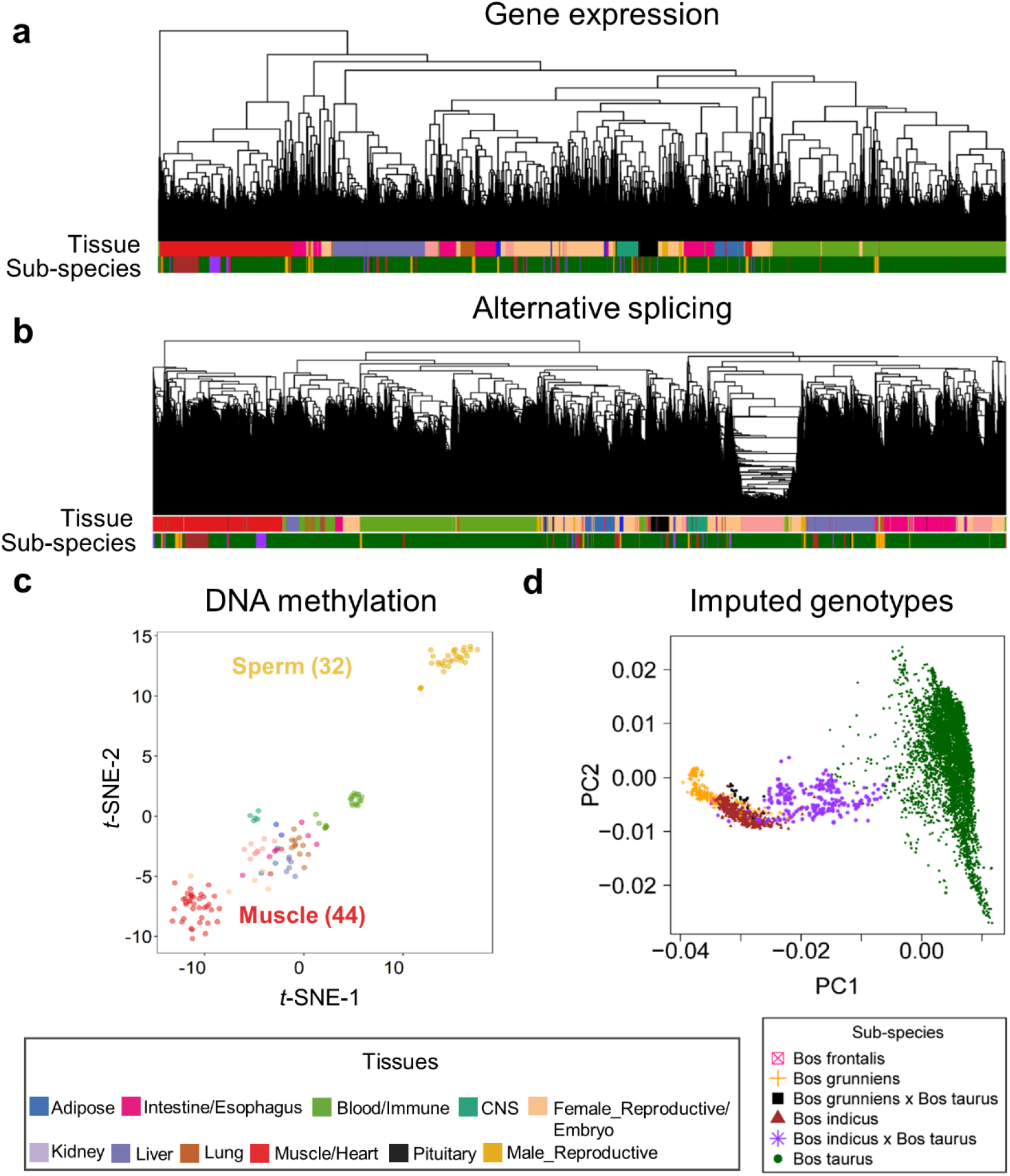
Hierarchical clustering and principal component analysis of samples. **(a)** Sample (n = 7,180) hierarchical clustering based on expression levels of all transcribed genes (Transcripts Per Million, TPM > 0.1). **(b)** Sample (7,180) hierarchical clustering based on alternative splicing value (Percent Spliced-In, PSI) of spliced introns. **(c)** Sample (n = 144) clustering using *t*-distributed SNE coordinates based on DNA methylation levels of CpG sites (coverage ≥ 5×). **(d)** Principal component analysis of samples (n = 7,180) based on imputed genotypes.

### Tissue specificity of transcriptome and methylome

Tissue-specificity of gene expression was significantly conserved between cattle and humans (Fig. 2a), and the function of genes with tissue-specific expression accurately reflected the known biology of the tissues. For instance, brain-specific genes were significantly enriched for synapse and neuron function, and testis-specific genes for spermatogenesis and reproduction (Fig. S5a). We also calculated tissue-specificity of promoter DNA methylation and gene alternative splicing. Similarly, the function of genes with tissue-specific promoter hypomethylation and splicing reflected the known biology of the tissues (Fig. S5b-c). We found that, based on tissue-specificity, the gene expression level was significantly and negatively correlated with DNA methylation level in promoters (Fig. 2b), and positively correlated with splicing ratios of introns (Fig. 2c). For example, *CELF2*, a brain-related gene, had a significantly higher expression, lower promoter DNA methylation, and higher splicing ratio of first intron in brain than in other tissues considered (Fig. 2d). Tissue-specific genes exhibited distinct patterns of sequence constraints (Fig. S5d), supporting the hypothesis of tissue-driven genome evolution^4^. We found that while brain-specific genes evolve slowly, blood or testis-specific ones evolve rapidly. This trend was also observed within tissue-specific hypomethylated regions (Fig. S5e-f).

**Fig. 2.**
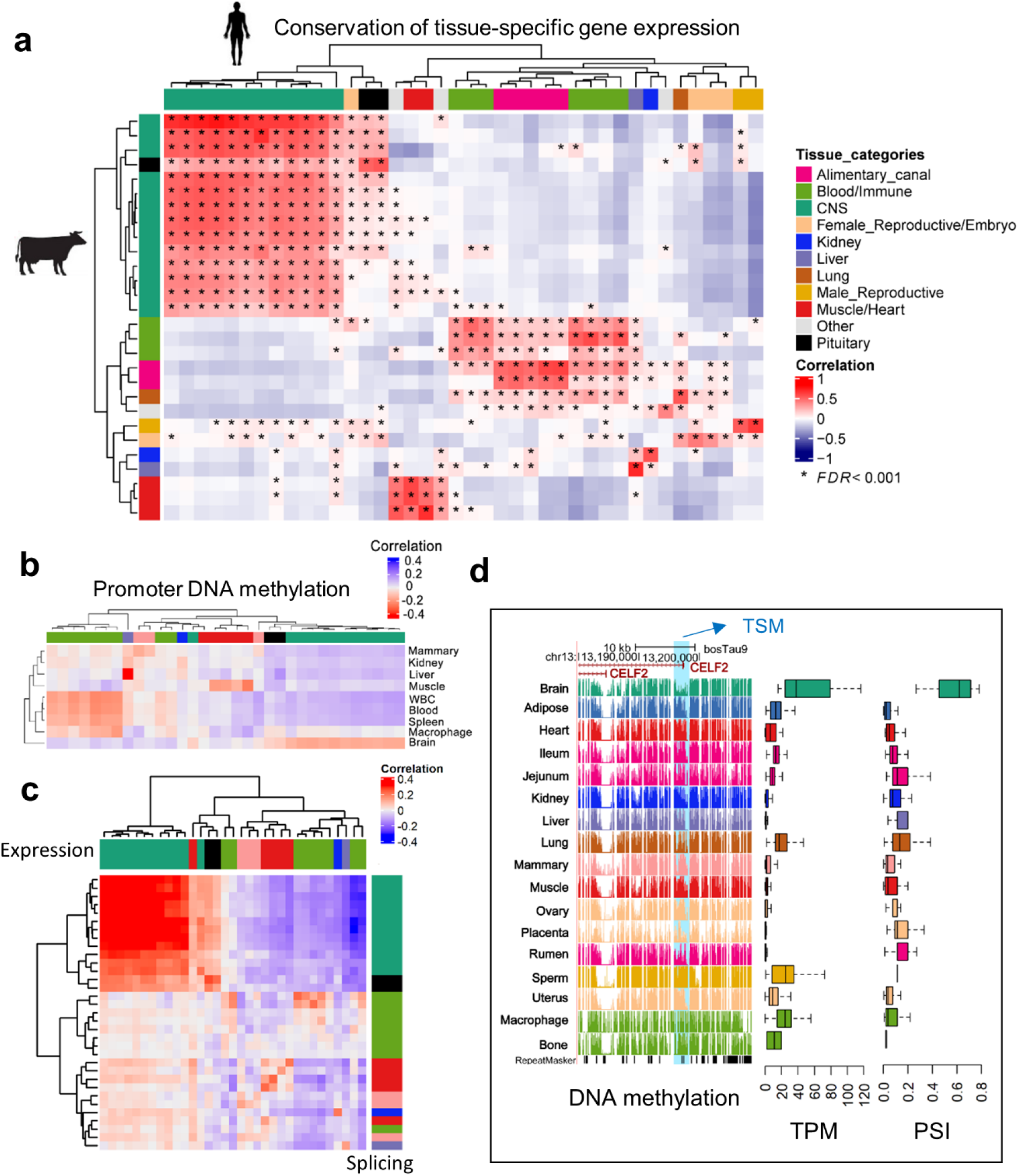
Tissue-specificity of gene expression, alternative splicing and DNA methylation. **(a)** Pearson correlation of tissue-specificity (measured as *t*-statistics) of 22,752 orthologous genes between cattle and humans tissues (GTEx v8)^3^. The multiple testing is corrected for using FDR. **(b)** Pearson correlation of tissue-specificity between gene expression (*x*-axis) and promoter DNA methylation levels (*y*-axis). WBC is for white blood cells. The color code of tissues in *x*-axis is the same as that in (**a). (c)** Pearson correlation of tissue-specificity between gene expression (Transcripts Per Million, TPM, *x*-axis) and alternative splicing (Percent Spliced-In, PSI, *y*-axis). The color code of tissues is the same as that in (**a). (d)** *CELF2* shows higher expression, lower DNA methylation levels in splice sites and higher PSI value of spliced introns (chr13:13034717−13197300) in brain tissue compared to the rest of tissues. TSM is for tissue-specific methylation.

### Discovery of expression and splicing QTLs

We identified *cis*-e/sQTLs for 24 tissues with 40 or more individuals, while accounting for relevant confounding factors and multiple testing (Fig. S6a-b). The number of eGenes (genes with significant *cis*-eQTLs) discovered ranged from 172 in ileum to 10,157 in blood, with 19,559 (83% of all 23,523 tested genes) classed as eGenes in at least one tissue (Table S4). The number of sGenes (genes with significant *cis*-sQTLs) discovered ranged from four in the salivary gland to 7,913 in macrophages, with 15,376 (70.8%) classed as sGenes in at least one tissue. Genes with no *cis*-eQTLs or -sQTLs (non-e/sGenes) in any of the tissues were significantly enriched in hormone activity, regulation of receptor activity, neuropeptide signaling pathway, and reproduction (Table S5). In general, the larger the number of samples for the tissue, the larger the number of *cis*-e/sGenes detected (Fig. 3a-b). As expected, with a larger sample size, we had more power to detect *cis*-eQTLs with smaller effect sizes (Fig. S7a-b). Consistent with findings in humans^13^, significant variants (eVariants) centered around the transcript start sites (TSS) of the measured genes (Fig. S7c-d). Across 24 tissues, an average of 46% (range 25.5 - 76.6%) of eVariants were found within 100 kb of the TSS of the target genes. In non-eGenes, there was also an enrichment of SNPs with the smallest *P*-values (but not statistically significant at FDR of 0.05) around TSS, suggesting a lack of power to detect such associations for those genes (Fig. S7c). Furthermore, we fine-mapped eGenes to assess whether the identified signals could be attributed to one or more causal SNPs. We found that an average of 46% (range 14.5 - 73.9%) of eGenes across 24 tissues had more than one independent *cis*-eQTLs (Fig. 3c), indicating the complex genetic control of gene expression. SNPs with the larger effects within a locus tended to be closer to the TSS (Fig. 3d). To complement and validate the *cis*-eQTL analysis within individuals, we conducted an allele-specific expression (ASE) analysis, and found that *cis*-eQTLs were significantly overrepresented in loci with significant (FDR < 0.05) ASE (Fig. 3e), and the effect sizes of *cis*-eQTLs was significantly correlated with those of ASEs (Fig. 3f, Fig. S7e-f).

**Fig. 3.**
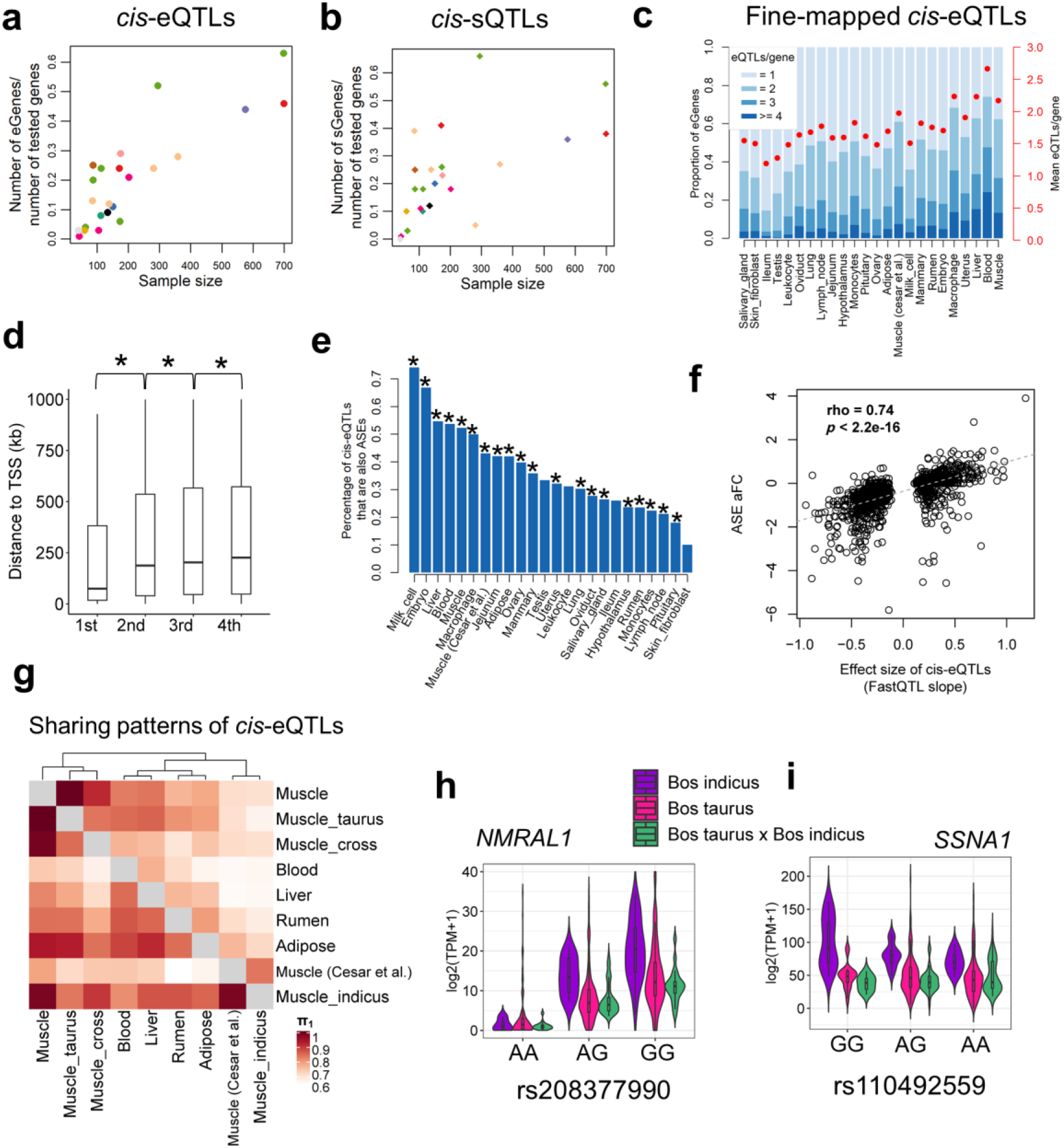
Discovery and characterization of *cis*-eQTLs and *cis*-sQTLs. **(a)** Relationship between the percentage of eGenes over all tested genes and sample size (Pearson *r* = 0.85; *P* = 1.30×10^−7^) across 24 tissues. **(b)** Relationship between the percentage of sGenes over all tested genes and sample size (Pearson *r* = 0.63; *P* = 1.06×10^−3^). **(c)** Distribution and average number of conditionally independent eQTLs per gene across tissues. Tissues are ordered by sample size. **(d)** The distance to transcription start site (TSS) increases from the 1^st^ to 4^th^ independent eQTLs. * indicates *P* < 0.05 based on the Student *t*-test. **(e)** *cis*-eQTLs are significantly (*P* < 1×10^−14^, denoted as *, Fisher Exact test) overrepresented in the loci with allelic specific expression (ASE). The *y*-axis indicates the percentage of *cis*-eQTLs that are also ASEs over all tested SNPs in the ASE analysis. **(f)** Correlation of effect sizes (FastQTL slope) of *cis*-eQTLs and allelic fold change (aFC) of ASEs (Spearman’s rho = 0.74, *P* < 2.2×10^−16^) in liver. (**g**) Pairwise *cis*-eQTL sharing patterns (π_1_ value) of muscle tissue across three subspecies (*Bos indicus, Bos taurus* and their crosses) and other tissues. Rows are discovery tissues, while columns are validation tissues. Muscle (Cesar et al.) is for 160 skeletal muscle samples of *Bos indicus* downloaded from Cesar et al. 2018^9^. **(h)** A *cis*-eQTL (rs208377990) of *NMRAL1* in muscle is shared across *Bos indicus, Bos taurus* and their crosses. **(i)** A *cis*-eQTL (rs110492559) of *SSNA1* in muscle is specific in *Bos indicus* (MAF = 0.25 and 0.37 in *Bos taurus* and *Bos indicus*, respectively), and has a significant (*p* < 5.61×10^−3^) genotype × subspecies interaction.

To investigate whether eQTLs are conserved among sub-species, we conducted *cis*-eQTL mapping for muscle samples from *Bos indicus* (n = 51 and 160), *Bos taurus* (n = 505), and their hybrids (n = 108) separately, yielding 86 (3,521), 2,766, and 800 eGenes, respectively. We observed that *cis*-eQTLs were more conserved across sub-species than across tissues (Fig. 3g). For example, the expression of *NMRAL1* in muscle was consistently and significantly regulated by a *cis*-eQTL (rs208377990) among *Bos indicus, Bos taurus*, and their hybrids (Fig. 3h). Combining the summary statistics of each sub-species in a meta-analysis showed that eGene-eVariant associations identified in one sub-species are potentially transferable to other sub-species (Fig. S7g-h). Combining samples from different sub-species and breeds will increase statistical power for detecting shared eQTLs, and enable more accurate mapping of the causal variants *via* reducing the linkage disequilibrium (LD) patterns. In total, 131 out of 437 eGene-eVariant pairs that were specifically discovered in *Bos indicus* showed significant (FDR < 0.05) genotype × subspecies interactions (Table S6). For instance, the expression of an immune-related gene, *SSNA1*, was regulated by a *cis*-eQTL (rs110492559) in *Bos indicus* but not in *Bos taurus* or the hybrids, showing a significant (*p* < 5.61×10^−3^) genotype × subspecies interaction (Fig. 3i). In addition, we found that subspecies-specific eQTLs had lower minor allele frequency (MAF) than subspecies-common eQTLs, consistent in both *Bos indicus* and *Bos taurus* (Fig. S8). This may indicate that the difference in eQTLs between subspecies could be partially due to their difference in the frequency of the segregating variants, provided that there are no epistatic/environmental/developmental effects.

The tissue-sharing patterns of *cis*-QTLs could provide novel insights into molecular regulatory mechanisms underlying complex phenotypes^3^. We applied the π_1_ statistics to measure the sharing patterns of *cis*-e/sQTLs between tissues (Fig. 4a and Fig. S9a). In general, we observed that both *cis*-eQTLs and *cis*-sQTLs tended to be tissue-specific or ubiquitous across tissues (Fig. 4b). We also calculated the tissue-sharing patterns of gene expression and alternative splicing (Fig. S9b-c), and found that the tissue-sharing patterns of the four core data types (i.e., gene expression, alternative splicing and cis-e/sQTLs) were significantly correlated (Fig. 4c). This result suggests that tissues with similar transcriptional profiles shared the genetic regulatory mechanisms of transcription. Further analysis on the expression of eGenes across tissues revealed that effect sizes of eVariants decreased with the increasing number of tissues where target eGenes were expressed (Pearson’s *r* = -0.27, *P* < 2.2×10^−16^), indicating that, on average, tissue-specific genes might be regulated by SNPs with larger genetic regulatory effects than widely-expressed genes (Fig. 4d). Due to limitations and challenges of *trans*-eQTLs analysis in this study which include: insufficient statistical power, the relatively lower imputation accuracy of distant intergenic SNPs, and complex inter-chromosomal LD in cattle (which could lead to increased type I error rates)^14^, we only conducted an exploratory *trans*-e/sQTL mapping for 15 tissues with over 100 individuals. We detected an average of 1,058 and 84 *trans*-eGenes and *trans*-sGenes (FDR < 0.05) across tissues, respectively (Table S7). We summarized the details of *trans*-eQTL mapping, including LD patterns of *trans*-eQTLs and *cis*-eQTL, tissue-sharing patterns of *trans*-eQTLs and their validations, in Fig. S10-11.

**Fig. 4.**
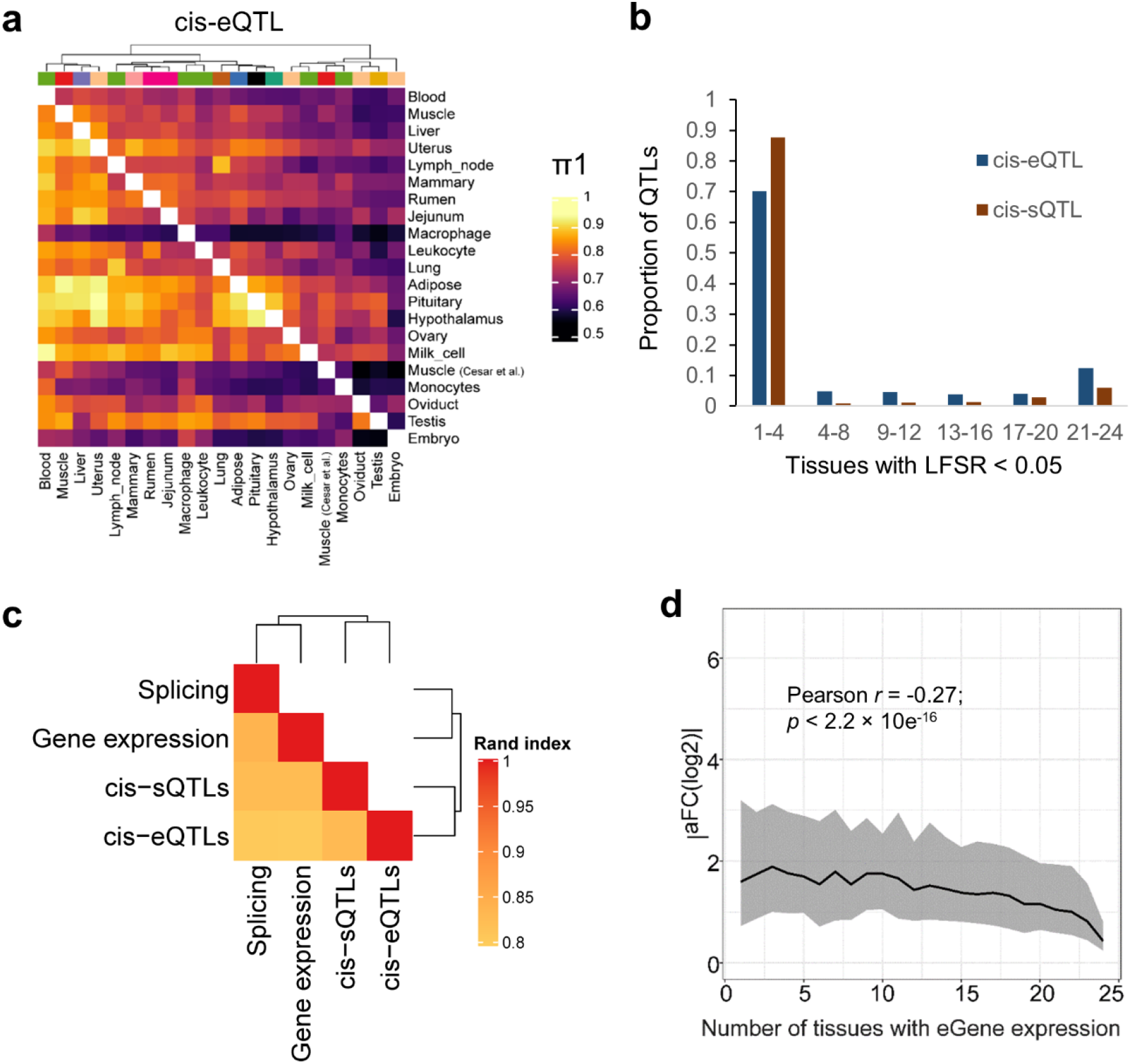
Tissue-sharing patterns of *cis*-QTLs. **(a)** Pairwise *cis*-eQTL sharing patterns (π_1_ value) across 24 tissues. **(b)** Tissue activity of *cis*-eQTLs and *cis*-sQTLs, where a *cis*-QTL is considered active in a tissue if it has a *mashr* local false sign rate (LFSR, equivalent to FDR) of < 5%. **(c)** The similarity of tissue clustering across four data types (*cis*-eQTL, *cis*-sQTL, gene expression and splicing)^3^. The k-means clustering, implemented in the fossil v0.4.0 R package^53^, is performed based on 2-22 clusters with 100,000 iterations. For each pairwise data types, we report the median Pairwise Rand index across all clusters. **(d)** Median (line) and interquartile range (shading) of *cis*-eQTL effect size (*y*-axis, measured as the absolute log_2_ transformed allele Fold Change, |aFC(log2)|), as a function of the number of tissues in which the eGene is expressed (*x*-axis; TPM > 0.1). Pearson correlation between |aFC(log2)| and number of tissues with eGene expression is −0.27, with *p* value < 2.2×10^−16^.

### Functional annotation of QTLs

We employed multiple layers of biological data to better define the molecular mechanisms of genetic regulatory effects. As expected, *cis*-e/sQTLs were significantly enriched in functional elements, such as 3’UTR and open chromatin regions (defined by ATAC-Seq data in cattle rumen epithelial primary cells)^15^ (Fig. 5a-b). Similarly, *cis*-sQTLs had a higher enrichment in splice donors/acceptors than *cis*-eQTLs. The *cis*-eQTLs associated with stop gains had larger effect sizes than other *cis*-eQTLs (Fig. 5c). The *cis*-e/sQTLs were enriched in hypomethylated regions of the matching tissues across 13 tissues (Fig. 5d-e). For instance, the liver exhibited the highest enrichment of *cis*-e/sQTL in liver-specific hypomethylated regions. Consistent with the brain having distinct abundance of alternative splicing, related to the development of the nervous system^13^, *cis*-sQTLs in the hypothalamus and pituitary had the highest enrichments in their specific hypomethylated regions (Fig. 5e).

**Fig. 5.**
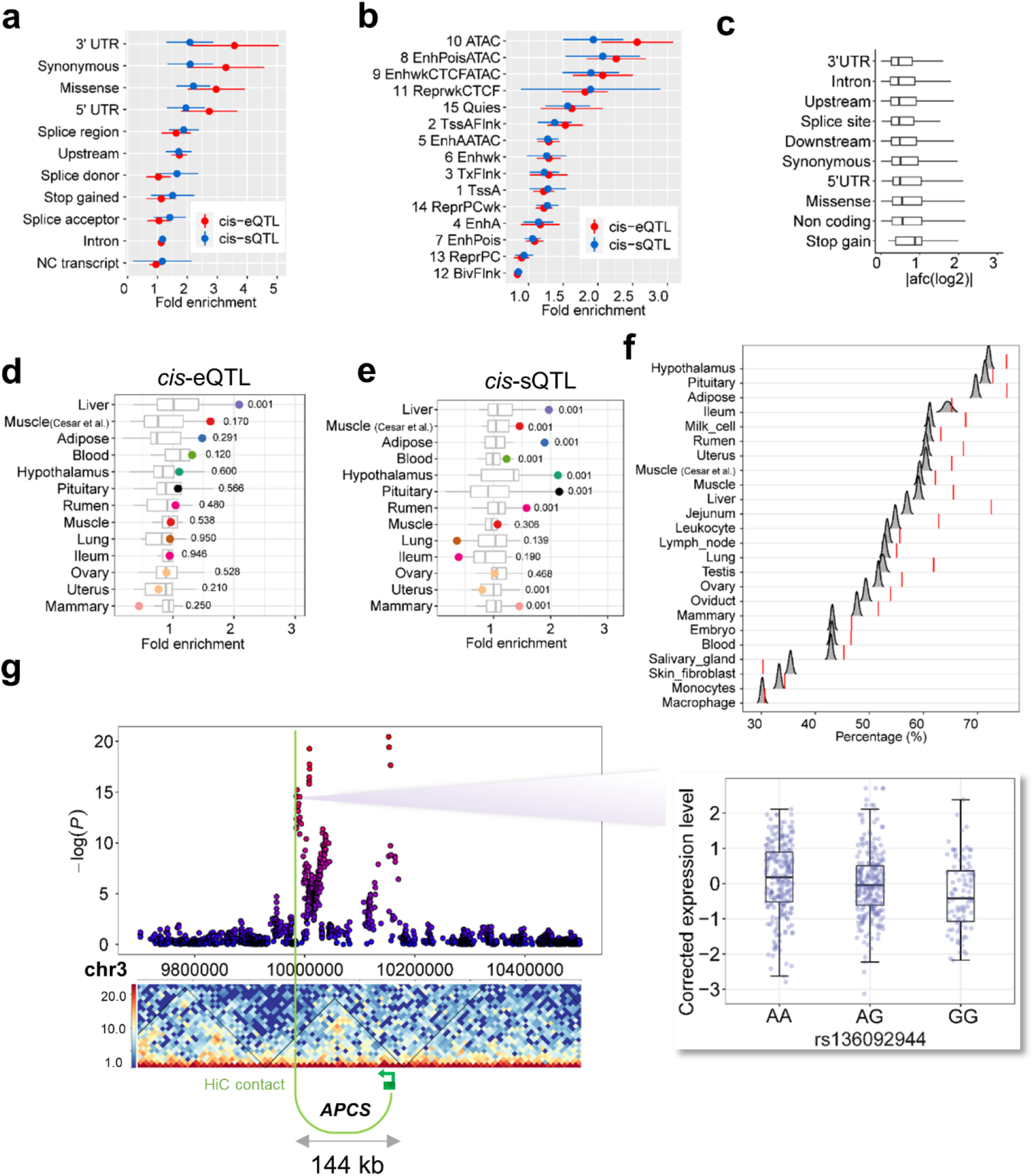
Functional annotation of *cis*-QTLs. **(a)** Enrichment of *cis*-eQTLs and *cis*-sQTLs of 24 tissues in sequence ontology. **(b)** Enrichment of *cis*-eQTLs and *cis*-sQTLs of 24 tissues in 15 chromatin states predicted from cattle rumen epithelial primary cells in Holstein animals^14^. **(c)** Effect sizes (measured as |aFC(log_2_|) of *cis*-eQTLs of 24 tissues across sequence ontology. **(d)** and **(e)** Enrichment of *cis*-eQTLs and *cis*-sQTLs of 13 tissues in tissue-specific hypomethylated regions, respectively. These 13 tissues have both DNA methylation and *cis*-QTL data. Higher fold enrichments are observed for matched tissues (highlighted dots) compared to other tissues. The numbers are *p*-values for enrichments of matched tissues based on the permutation test (times = 1,000). **(f)** Percentages of eGene-eVariant pairs that are located within topologically associating domains (TADs) are significantly (FDR < 0.01) higher than those of random eGene-SNP pairs with matched distance, except for ileum, macrophage and skin fibroblast. The TADs are obtained from the lung Hi-C data. The null distributions of percentages of eGene-SNP pairs within TADs are obtained by doing 5,000 bootstraps. **(g)** An eGene (*APCS*) and its eVariant (rs136092944) are located within a TAD, and linked by a significant Hi-C contact in cattle lung tissue. The Manhattan plot shows the *P*-values of all tested SNPs in the *cis*-eQTL mapping analysis of *APCS*. The boxplot (right) shows the PEER-corrected expression levels of *APCS* across the three genotypes of eVariant (rs136092944), i.e., AA, AG, and GG, respectively.

Topologically associated domains (TADs) enable chromatin interactions between distant regulatory regions and target promoters^16^. By examining Hi-C data of lung tissue in cattle^15^, we obtained TADs and significant Hi-C contacts, which were likely to be conserved across tissues as proposed previously^16^. By comparing with random eGene-SNP pairs with matched distances, we observed significantly (FDR < 0.01, 5,000 bootstrapping test) higher percentages of eGene-eVariant pairs within TADs across the majority of tissues, except for ileum and skin fibroblast (Fig. 5f). For instance, *APCS* and its *cis*-eQTL peak (144kb upstream of the TSS) were encompassed by an TAD and linked by a significant Hi-C contact, which allowed the regulation of its expression by a distant eVariant (rs136092944) (Fig. 5g).

### eQTLs and complex trait associations

The primary goal of this study is to provide a resource for elucidating the genetic and biological mechanisms involved in cattle. We thus evaluated e/sQTLs detected in each tissue for associations with four distinct agronomic traits, i.e., ketosis, somatic cell score in milk (SCS), age at first calving (AFC), and milk yield (MY). The top SNPs associated with ketosis from GWAS were significantly (*P* < 0.05, 1,000 permutation test) enriched within liver *cis*-e/sQTLs (Fig. 6a). Similarly, MY-associated SNPs were significantly overrepresented in mammary gland *cis*-e/sQTLs (Fig. 6b). Compared to other tissues, mammary gland, milk cells and liver were the tissues with highest enrichment of MY-associated SNPs amongst *cis*-eQTLs (Fig. 6c). Additionally, AFC-associated SNPs were significantly enriched for monocytes *cis*-eQTLs, and SCS for mammary gland (Fig. S12a-b).

**Fig. 6.**
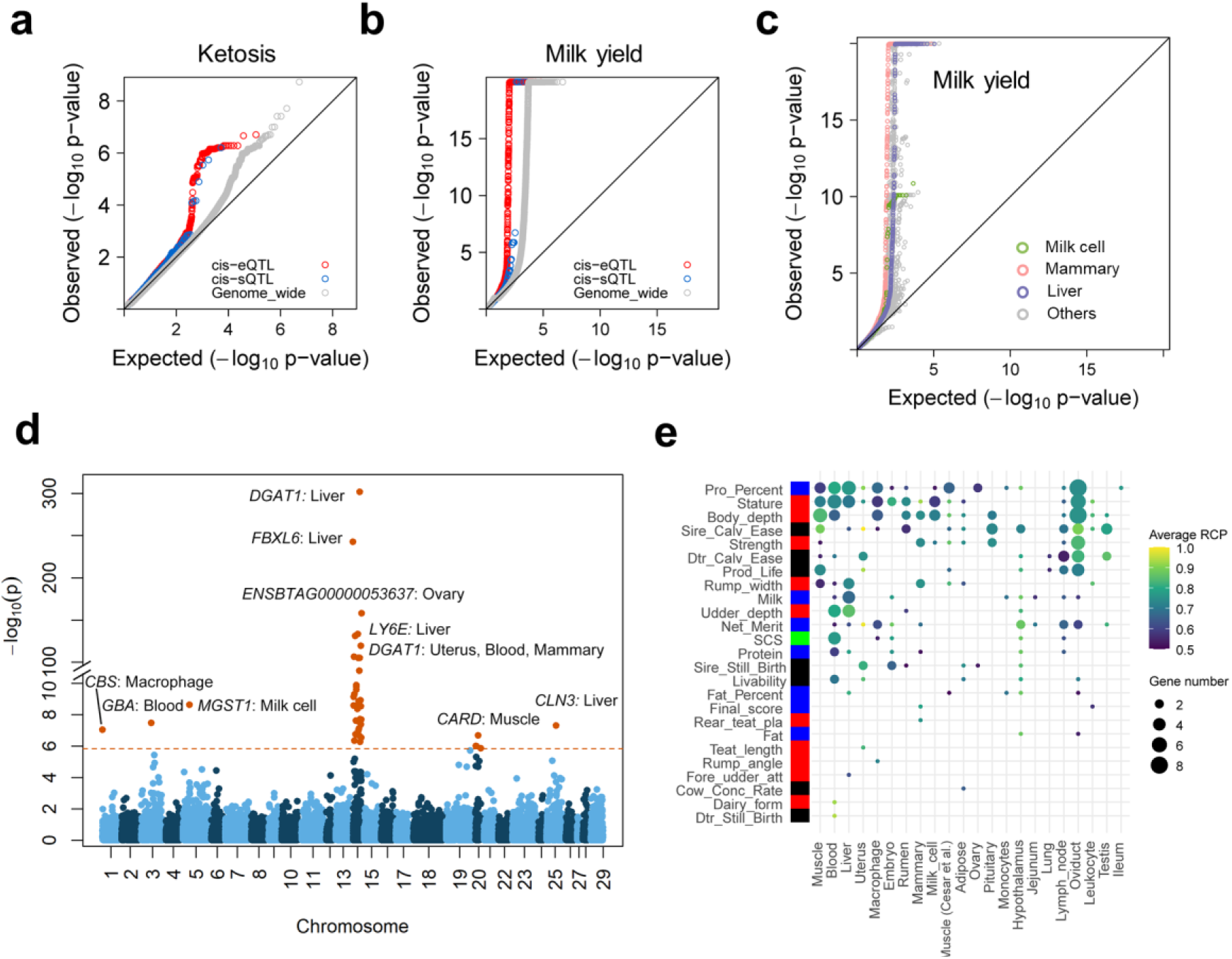
Relationship between complex traits and *cis*-QTLs. **(a)** *cis*-eQTLs (*P* = 0.001, 1,000 permutations) and *cis*-sQTLs (*P* = 0.02) in liver show significantly higher enrichments for top SNPs associated with ketosis compared to genome-wide SNPs (shown in grey). **(b)** *cis*-eQTLs (*P* = 0.001) and *cis*-sQTLs (*P* = 0.03) in mammary gland show higher enrichments for top SNPs associated with milk yield compared to genome-wide SNPs (shown in grey). **(c)** Enrichment of *cis*-eQTLs for genetic associations with milk yield is tissue-dependent. The *cis*-eQTLs in mammary gland, milk cells and liver exhibit higher enrichments for genetic associations with milk yield compared to those in other tissues. **(d)** Manhattan plots of transcriptome-wide association study (TWAS) for milk yield across all 24 tissues. **(e)** The number of genes that were colocalized (regional colocalization probability, *rcp* > 0.5 in fastENLOC) between GWAS significant loci of complex traits and cis-eQTLs across tissues. The size of point indicates the number of genes, while the color of point indicates the average *rcp* of each trait-tissue pair.

We detected 854 significant gene-trait pairs for 43 agronomic traits in cattle *via* single-tissue TWAS, representing 337 unique genes (Table S8). Out of 319 previously fine-mapped genes^17,18^, we validated 54, including linking expression of *DGAT1* in liver and mammary gland, and expression of *MGST1* in milk cells, as well as expression of *CLN3* in liver to MY (Fig. 6d). The expression of *ZNF613* in hypothalamus was the most significant association for many reproduction and body conformation traits, including daughter-still-birth and stature (Table S8), supporting our previous finding that *ZNF613* is significantly associated with gestation length possibly through its influence on embryonic development^19^. Furthermore, we conducted a colocalization analysis of *cis*-eQTLs and GWAS loci, and detected 115 unique eGenes that were colocalized (regional colocalization probability, *rcp* > 0.5) within 260 GWAS loci associated to 25 out of the 43 complex traits analysed. These represent 235 significant gene-trait pairs (some eGenes were associated with 2 or more traits) (Fig. 6e; Table S9). For instance, *TIGAR*, a muscle *cis*-eGene, playing roles in cellular metabolism and oxidative stress, was colocalized (*rcp* = 0.529) with a GWAS locus associated with strength on chromosome 5 (Fig. S12c-d). We also took sire calving ease, which GWAS loci were colocalized with 21 eGenes in at least one tissue, as an example in Fig. S12e. By comparing results from single/multi-tissue TWAS and colocalization, we found an overlap of 66 gene-trait pairs (Table S10; Fig. S12f). Overall, TWAS and colocalization analyses enhanced our ability to detect candidate causal genes and to better understand the biological underpinnings of complex traits in cattle.

## Discussion

The cattle GTEx atlas represents the most comprehensive reference resource of the cattle transcriptome to date. It provides a detailed characterization of genetic control of gene expression and splicing across 24 tissues in cattle. This study demonstrates that it is possible to discover genetic regulatory variants of transcriptome by deriving and imputing genetic variants from RNA-Seq data only in livestock. We established a *in silico* protocol to deliver a livestock GTEx atlas in a timely manner and at a fraction of the cost of the human GTEx project, or an equivalent project in livestock generating RNA-Seq data from scratch. Although we have successfully uncovered large numbers of tissue-specific e/sQTLs and provided a comprehensive view of the control of gene regulation in cattle and an accompanying public database that is a valuable resource for the community, we are also mindful that this resource can be further improved with the inclusion of more individuals/breeds and more varied data types. The imputation accuracy for breeds that are very under-represented in the reference panel might be relatively low. Additionally, generating SNP array genotypes or WGS for individuals with RNA-Seq data can provide additional information for distant intergenic variants as compared to RNA-Seq data only, potentially enhancing the detection of distant intergenic QTLs. The Farm animal GTEx (FarmGTEx) consortium was recently launched and is currently extending the bioinformatics pipeline developed here to other livestock species (e.g., pigs, small ruminants and chicken) to add value to the publicly available sequencing data for the research community.

The cattle GTEx provides a resource to explore tissue-sharing patterns of the transcriptome and its genetic regulation (i.e., e/sQTLs) in cattle. In contrast to the human GTEx^4^, where RNA-Seq samples across tissues were collected from the same individuals, the cattle GTEx used publicly available data, where individuals or even breeds were different from tissue to tissue. This might explain why there is a lower proportion of *cis*-eQTLs and *cis*-sQTLs shared across tissues compared to the human GTEx. In addition, the difference in the cell type composition of tissues can also affect the tissue-sharing patterns of QTLs^4^. When single-cell RNA-Seq data is available for multiple tissues in cattle in the near future^20^, it would be of interest to computationally estimate the cell type proportions in the bulk tissue samples to uncover the cellular specificity of genetic regulatory effects^21^.

This cattle GTEx atlas systematically links SNPs, genes, and tissues for the first time in cattle, and provides an important tool for new discoveries using these three datasets to study the mechanisms underlying complex traits. The e/sQTLs detected here provide a rich set of functional variants for agronomic traits in cattle, as we found that top GWAS associations of traits were significantly enriched for regulatory QTLs in their relevant tissues. Our TWAS and colocalization analyses further provide a list of promising candidate genes/variants for functional follow-up. We noted the relatively small overlap of results from TWAS and colocalization. This might because the two methods use the information differently, with the TWAS or co-localization method being more or less powerful depending on the genetic architecture of both the trait of interest and the tissue gene expression.

Further integration of these QTLs with functional annotations of a range of tissues from the on-going Functional Annotation of Animal Genomes (FAANG) project will provide valuable opportunities to understand transcriptional/post-transcriptional regulatory mechanisms underpinning GWAS hits for agronomic traits^22^. The multi-tissue e/sQTLs generated here will also enable the exploration of molecular mechanisms underlying the extensive pleiotropic effects identified in livestock^23^. This information will allow the understanding of mechanisms of response to intended selection as well as disentangling unintended and unfavorable correlated responses to this same selection (e.g. increasing mastitis or deteriorating fertility when selection for increased milk production). Furthermore, this resource will assist in the development of genomic selection methods and tools to improve animal health and wellbeing. For instance, a better understanding of the genetic architecture underpinning agronomic traits will benefit genetic improvement programs by incorporating biological knowledge into genomic prediction models^10^, which has been shown to improve prediction accuracy across generations, populations and breeds^23^.

## Online Methods

### Quantification of gene expression

We downloaded 11,642 RNA-Seq datasets (by July, 2019) from SRA (n = 11,513, https://www.ncbi.nlm.nih.gov/sra/) and BIGD databases (n = 129, https://bigd.big.ac.cn/bioproject/). We merged multiple datasets from single samples, yielding 8,536 unique RNA-Seq samples. We applied a stringent and uniform pipeline to filter and analyze all the data. Briefly, we first removed adaptors and low quality reads using Trimmomatic (v0.39)^24^ with parameters: adapters/TruSeq3-SE.fa:2:30:10 LEADING:3 TRAILING:3 SLIDINGWINDOW:4:15 MINLEN:36. We filtered out samples with clean read counts ≤ 500K, resulting in 7,680 samples, and mapped clean reads to the ARS-UCD1.2 cattle reference genome^25^ using single or paired mapping modules of STAR (v2.7.0) with parameters of outFilterMismatchNmax 3, outFilterMultimapNmax 10 and outFilterScoreMinOverLread 0.66. We kept 7,264 samples with uniquely mapping rates ≥ 60% (mean, 91.07%; range, 60.44%-100%; mapping details in Table S1). We then obtained normalized expression (TPM) of 27,608 Ensembl (v96) annotated genes using Stringtie (v2.1.1)^26^, and extracted raw read counts of them with featureCounts (v1.5.2)^27^. We finally clustered 7,264 samples based on log_2_(TPM +1) using a hierarchical clustering method, implemented in R package *dendextend*, with distance = (1-*r*), where *r* is the Pearson correlation coefficient. We excluded samples with obvious clustering errors (e.g., samples labeled as liver that were not clustered with other liver samples), resulting in 7,180 samples for subsequent analysis.

### Quantification of alternative splicing

We used Leafcutter (v0.2.9)^28^ to identify and quantify variable alternative splicing events of genes by leveraging information of junction reads (i.e., reads spanning introns) that were obtained from the STAR alignment. The Leafcutter enables the identification of splicing events without relying on existing annotations that are typically incomplete, especially in the setting of large genes or individual- and/or population-specific isoforms^28^. We first converted bam files from STAR alignment into junction files using the script “bam2junc.sh”, and then performed intron clustering using the script “leafcutter_cluster.py” with default settings of 50 reads per cluster and a maximum intron length of 500 kb. We employed the “prepare_genotype_table.py” script in Leafcutter to calculate intron excision ratios and to remove introns used in less than 40% of individuals or with no variation. Ultimately, we standardized and quantile normalized intron excision ratios as Percent Spliced-In (PSI) values across samples. We clustered 7,180 samples based on PSI using the same method as used in gene expression.

### Genotyping and imputation

We called genotypes of known genomic variants in the 1000 Bull Genomes Projects^10^ for 7,180 high-quality RNA-Seq samples individually, following the recommended best practices pipeline in Genome Analysis Toolkit (GATK) (v4.0.8.1)^29^ with default settings. We filtered out low quality SNPs using --filter-expression “FS > 30.0 & QD < 2.0”. We then imputed the filtered SNPs on autosomes to sequence level using Beagle (v5.1)^30^ based on a multiple-breed reference population consisted of 3,103 individuals from run7 of the 1000 Bull Genomes Project^10^ and 207 public individuals from *Bos taurus* (n = 101), *Bos indicus* (zebu, n = 20), and *Bos grunniens* (yak, n = 86) (Table S11). Finally, we obtained 6,123 samples that were genotyped and imputed successfully. We filtered out variants with MAF < 0.05 and dosage R-squared (DR^2^) < 0.8, resulting in 3,824,444 SNPs used for QTL mapping. To evaluate the accuracy of imputation, we called genotypes (∼6 M SNPs) from WGS (average read depth > 10×) of Holstein (n = 4), Limousin (n = 3) and Angus (n = 5) animals, which had RNA-Seq data as well. We then measured the genotype concordance rates between WGS-SNPs and RNA-Seq/imputed SNPs. We extracted 153,913 LD-independent SNPs using plink (v1.90)^31^ (--indep-pairwise 1000 5 0.2), and conducted PCA analysis for all 6,123 samples using these SNPs in EIGENSOFT (v7.2.1)^32^. We calculated the identity-by-state (IBS) distance among samples by using these independent SNPs to remove duplicate individuals. IBS distance = (IBS2 + 0.5*IBS1) / (IBS0 + IBS1 + IBS2), where IBS0 is the number of IBS 0 non-missing variants, IBS1 is the number of IBS 1 non-missing variants and IBS2 is the number of IBS 2 non-missing variants. We set an IBS distance cutoff of 0.85 to deem two samples as duplicates and kept one of them. When conducting QTL mapping, we removed an average of 43 duplicate samples within each tested tissue (ranging from one in salivary gland and leukocyte to 132 in muscle), resulting in 4,889 samples.

### Allele specific expression (ASE)

We conducted ASE analysis using the GATK ASEReadCounter tool (v4.0.8.1) with the following settings: --U ALLOW_N_CIGAR_READS -minDepth 10 –minMappingQuality 255 --minBaseQuality 10. SNPs for ASE detection fulfilled the following criteria: heterozygous in at least five samples, at least 10 reads per allele, and at least 2% of all reads supporting the minor allele. We then calculated a binominal *P*-value by comparing to the expected ratio under the null hypothesis, followed by multiple-test correction with the Benjamini–Hochberg approach (FDR). SNPs with FDR < 0.05 were considered as significant ASE. We estimated the effect size (allele fold change, aFC) of regulatory variants at ASE loci using a haplotype-based approach implemented in phASER^33^.

### Bioinformatics analysis of WGBS data

For WGBS data analysis, we first used FastQC (v0.11.2) and Trim Galore v0.4.0 (-- max_n 15 --quality 20 --length 20 -e 0.1) to determine read quality and to filter reads with low quality, respectively. We then mapped clean reads to the same reference genome (ARS-UCD1.2) using Bismark software (v0.14.5)^34^ with default parameters. After deduplication of reads, we extracted methylation levels of cytosines using the *bismark_methylation_extractor* (--ignore_r2 6) function. The coverages of all WGBS data were calculated using clean reads with an average of 27.6-fold coverage (range: 5-47 ×). Ultimately, we kept CpG sites that were represented by at least five reads for subsequent analyses. We visualized sample clusters based on DNA methylation levels of shared CpGs using *t*-SNE approaches.

### Identification of TAD and significant Hi-C contacts

To find potential chromatin interactions between distant eVariants and target eGenes, we identified TADs and Hi-C contacts from Hi-C data from lung tissue in cattle that was retrieved from NCBI Sequence Read Archive (SRA) under accessions: SRR5753600, SRR5753603, and SRR5753606. We used Trim Galore (v0.4.0) to trim adapter sequences and low-quality reads (--max_n 15 --quality 20 --length 20 -e 0.1), resulting in ∼820 million clean reads. We then mapped clean reads to the reference genome (ARS-UCD1.2) using BWA^35^. We applied HiCExplorer v3.4.1^36^ to build a Hi-C contact matrix with 10kb resolution and identified TAD with hicFindTAD. We kept TADs with FDR less than 0.01 to link eQTLs to eGenes. We further employed HiC-Pro (v2.11.4)^37^ to call Hi-C contacts with 10 kb resolution from Hi-C data. Briefly, HiC-Pro aligned clean reads to the reference genome with Bowtie2 (v2.3.5)^35^. After building a contact matrix, HiC-Pro generated intra- and inter-chromosomal maps and normalized them using the ICE normalization algorithm. We considered Hi-C contacts with FDR < 0.05 as significant.

### Tissue-specificity analysis of gene expression, alternative splicing and DNA methylation

To quantify tissue-specific expression of genes, we computed a *t*-statistics for each gene in each of the 114 tissues. We grouped 114 tissues into 13 categories (Table S1). We scaled the log_2_-transformed expression (i.e., log_2_TPM) of genes to have a mean of zero and variance of one within each tissue. We then fitted a linear model as described in^15^ for each gene in each tissue using the least squares method. When constructing the matrix of dummy variables (i.e., design matrix) for tissues, we denoted samples of the target tissue/cell type (e.g., CD4 cells) as ‘1’, while samples outside the target category (e.g., non-blood/immune tissues) as ‘-1’. We excluded samples within the same category (e.g., CD8 cells and lymphocytes) to detect genes with specific expression in each particular category, even if they were not specific to the target tissue within this category. We obtained *t*-statistics for each gene to measure its expression specificity in a given tissue. We considered the top 5% of genes ranked by largest *t*-statistics as genes with high tissue-specific expression. In order to explore the conservation of tissue-specific expression between cattle and humans, we employed the same method to quantify the tissue-specific expression of all orthologous genes in each of 55 human tissues using GTEx (v8) data^3^.

To detect tissue-specific alternative splicing, we used leafcutter to analyze the differential intron excision by comparing the samples from the target tissue to the remaining tissues^28^, while excluding samples from tissues of the same category as the target tissue. We used the Benjamini-Hochberg method (FDR) to control multiple testing.

For DNA methylation, we focused on gene promoters (from upstream 1500bp to downstream 500bp of TSS), the methylation levels of which were calculated with a weighted methylation method using the roimethstat function in MethPipe (v3.4.3)^38^. We computed a *t*-statistic for the promoter of each gene using the same method as in tissue-specific expression analysis. We considered the bottom 5% of genes ranked by *t*-statistics as genes with tissue-specific promoter hypomethylation. We also detected tissue-specific methylation regions in a genome-wide mode using SMART2^39^ with parameters of -t DeNovoDMR -MR 0.5 -AG 1.0 -MS 0.5 -ED 0.2 -SM 0.6 -CD 500 -CN 5 -SL 20 -PD 0.05 -PM 0.05.

### Covariate analysis for QTL discovery

To account for hidden batch effects and other technical/biological sources of transcriptome-wide variation in gene expression, we estimated latent covariates in each tissue using the Probabilistic Estimation of Expression Residuals (PEER) method^40^. In each tissue, we estimated 75 PEER factors first. The posterior variances of factor weights dramatically decreased and reached or nearly reached plains when 10 PEER factors were included (Fig. S6a). Therefore, we used 10 PEER covariates to account for the effects of confounding variables on gene expression in all following QTL analyses. For instance, the variance of gene expression among samples in adipose captured by 9 out of 10 PEER factors were significantly (FDR < 0.05) correlated with known technical and biological covariates like clean data size, mapping rate, project, breeds, sub-species, sex and age (Fig. S6b). To further control the effect of population structure on the discovery of QTLs, we included genotype PCs based on sample size bins: three PCs for tissues with < 150 samples, five PCs for tissues with ≥ 150 and < 250 samples, and ten PCs for tissues with ≥ 250 samples.

### *cis*-eQTL mapping

We conducted *cis*-eQTL mapping for 24 tissues with at least 40 individuals each, while adjusting for corresponding PEER factors and genotype PCs. Detailed information about these 24 tissues is in Table S4. As the majority of *cis*-eQTLs are shared across sub-species/breeds (Fig. 3g), we combined, adjusting for species/breed, all of the datasets from the same tissue to perform *cis*-eQTL mapping in order to increase the statistical power. We kept genes with TPM > 0.1 in ≥ 20% samples in each tissue. Gene expression values of all samples in a given tissue were quantile normalized to the average empirical distribution and expression values for each gene then inverse normal transformed (INT) across samples. The *cis*-eQTL mapping was done using a linear regression model, implemented in FastQTL^41^, to test associations of the normalized expression level of genes with genetic variants in 1Mb of TSS of target genes. We only considered imputed variants with MAF > 0.05 and at least four minor alleles across samples within the target tissue. We first conducted *cis*-eQTL mapping in a permutation mode with the setting --permute 1000 10000, to identify genes with at least one significant *cis*-eQTL (eGene). We considered FDR ≤ 0.05 as significant, which was calculated with the Benjamini-Hochberg method based on the beta distribution-extrapolated empirical *P*-values from FastQTL. To identify a list of significant eGene-eVariant pairs, we applied the nominal mode in FastQTL. A genome-wide empirical *P*-value threshold *p*_*t*_ was defined as the empirical *P*-value of the gene closest to the 0.05 FDR threshold^3^. We then calculated the nominal threshold as *F*^−1^(*p*_*t*_), where *F*^−1^ is the binominal inverse cumulative distribution, of which parameters for genes were obtained from the above permutation mode of FastQTL analysis. We considered variants with nominal *P*-values below the nominal threshold as significant, and included them into the list of eGene-eVariant pairs. We calculated the aFC, defined as the ratio of the expression level of the haplotype carrying the alternative allele over the one carrying the reference allele, to measure effect sizes of *cis*-eQTLs using the aFC (v0.3) tools^42^. We further applied the statistical fine-mapping method, dap-g^43^, to infer multiple independent casual *cis*-eQTLs of a gene in a tissue. The dap-g approach employed a Bayesian variable selection model, using a signal-level posterior inclusion probability (SPIP) to measure the strength of each association signal (SNPs in LD). We set a cutoff of 0.1 (i.e., SPIP > 0.9) as the inclusion threshold to detect representative/independent eQTLs for the target eGene. To analyze pairwise tissue similarity in QTLs, we calculated π_1_ statistics, defined as the proportion of true positive QTLs identified in first tissue (Discovery tissue) amongst all tested gene-variant pairs in second tissue (Validation tissue), using the Storey and Tibshirani qvalue approach, as described in^13^.

### Meta-analysis of *cis*-eQTLs of muscle samples from three sub-species

Data from muscle samples were available from three sub-species: *Bos indicus* (n = 51), *Bos taurus* (n = 505), and their crosses (n = 108). To explore the similarity and variability of *cis*-eQTLs among sub-species, we conducted *cis*-eQTL mapping using muscle samples from each of the sub-species separately. We then conducted a meta-analysis to integrate *cis*-eQTL results from three sub-species using the METAL tool^44^. We obtained Z-scores (the sum of weighted effect sizes) of SNPs from the meta-analysis. Weights were proportional to the square-root of the number of individuals in each sub-species^44^. We employed plink^31^ (http://pngu.mgh.harvard.edu/purcell/plink/) to test the SNP × subspecies interaction in muscle samples, and adjusted the *p*-values to FDR using Benjamini-Hochberg procedure. We took *FDR* < 0.05 as the significant threshold.

### *cis*-sQTL mapping and tissue-sharing patterns

In each of the 24 tissues, we applied a linear regression model, implemented in FastQTL^41^, to test for associations of genotypes within 1 Mb up- and down-stream of target intron clusters and their corresponding intron excision ratios. We used the first five genotype PCs to account for the effect of ancestry, and 10 PEER factors to adjust for the effect of unknown confounding variables. We applied the permutation pass mode (--permute 1000 10000) in FastQTL^41^ to obtain beta approximated permutation *p* values, followed by multiple test correction with the FDR method. We considered sQTL-intron pairs with FDR < 0.05 as significant, and defined sGene as genes containing a significant sQTL in any introns. We employed MashR^45^ to analyze tissue-sharing patterns of QTLs as described previously in human GTEx^3^, and considered the local false sign rate (LFSR) < 0.05 as significant.

### *trans*-QTL mapping

We conducted *trans*-eQTLs for 15 tissues with at least 100 samples each. We filtered genomic variants using a more stringent threshold than *cis*-eQTL mapping to partially account for the reduction in statistical power. We obtained mappability of variants based on k-mer lengths of 36 and 75 following the procedure described in https://wiki.bits.vib.be/index.php/Create_a_mappability_track. We excluded any variants within repeats (Repeatmasker and simple repeats), and further removed variants with mappability < 1, based on k-mer length of 75. After filtering, we kept SNPs with MAF > 0.05 and at least 10 minor alleles within each tissue for association testing.

We applied two methods to detect *trans*-eQTLs for protein-coding genes with an average mappability ≥ 0.8 based on k-mer length of 36. Firstly, we associated the normalized expression of target genes with genotypes on other autosomal chromosomes using a linear regression model in MatrixQTL^46^, while adjusting for the same covariates as in *cis*-eQTL analysis. Secondly, we employed a linear mixed model (by fitting a polygenic effect with the genetic relationship matrix to further account for the complex relatedness among individuals) in the GCTA software^47^ for *trans*-eQTL and *trans*-sQTL mapping. For both methods, we adjusted *P*-values for multiple testing using the Benjamini-Hochberg method to obtain FDR. We considered gene-variant pairs with FDR < 0.05 as significant. To conduct an internal validation of *trans*-eQTL mapping, we randomly and evenly divided blood and muscle samples into two groups. We first conducted *trans*-eQTL mapping in the first group using the linear mixed model to detect significant *trans*-eQTL-gene pairs, and then repeated in the second group.

### TWAS and Colocalization of *cis*-eQTLs and GWAS loci

To associate gene expression in a tissue with complex traits, we conducted a single-tissue TWAS analysis using S-PrediXcan^48^ by prioritizing GWAS summary statistics for 43 agronomic traits of economic importance in cattle, including reproduction (n = 11), production (milk-relevant; n = 6), body type (n = 17), and health (immune/metabolic-relevant; n = 9). For body conformation (type), reproduction, and production traits, we conducted a single-marker GWAS by fitting a linear mixed model in 27,214 U.S. Holstein bulls as described previously^17^. For health traits, we conducted GWAS using the same method in a subset (ranging from 11,880 for hypocalcemia to 24,699 for livability) of the 27,214 available bulls^18^. We constructed a Nested Cross Validated Elastic Net prediction model using genotype and expression data. We included sub-species, 10 PEER factors and corresponding genotype PCs in the model to adjust for unknown confounding variables and underlying population structure. For each trait, we conducted TWAS in each of the same 24 tissues as in *cis*-eQTL mapping. We considered genes with Bonferroni-corrected *P* < 0.05 as significant. We visualized the Manhattan plots of *P*-values of all tested genes using ggplot2 (v3.3.2) in R (v3.4.1). In addition, we further employed S-MultiXcan^49^ to conduct multi-tissue TWAS analysis, and considered gene-trait pairs with Bonferroni threshold *p* < 4×10^−6^ (0.05/number of tested genes) significant.

To detect the shared causal variants of gene expression and complex traits, we conducted a colocalization analysis of *cis*-eQTLs from 24 tissues and GWAS loci of 43 agronomic traits using fastENLOC v1.0^50^. Briefly, we split the imputed GWAS summary statistics into approximately LD-independent regions, and each region was considered as a GWAS locus. The LD-independent regions were generated from genotypes of 886 Holstein animals from run7 of 1000 bull Genomes project, as the GWAS summary statistics were from the U.S. Holstein population. In each GWAS locus of a trait with suggestive significant SNPs (*P* < 10^− 5^), we considered a gene with regional colocalization probability (*rcp*) > 0.5 as significant.

### Other downstream bioinformatics analysis

We used Genomic Association Tester (GATv1.3.4)^51^ 1,000 permutations to estimate the functional enrichment of QTLs in particular genomic regions, e.g., chromatin states and methylation elements. We considered enrichments with FDR < 0.05 as significant. We used the R package, ClusterProfiler^52^, to annotate the function of genes based on the Gene Ontology database from Bioconductor (org.Bt.eg.db v3.11.4). We considered GO terms with FDR < 0.05 as significant.

## Acknowledgements

This work was supported in part by AFRI grant numbers 2016-67015-24886, 2019-67015-29321, and 2021-67015-33409 from the USDA National Institute of Food and Agriculture (NIFA) Animal Genome and Reproduction Programs and BARD grant number US-4997-17 from the US-Israel Binational Agricultural Research and Development (BARD) Fund. L. Fang. was partially funded through HDR-UK award HDR-9004 and the Marie Skłodowska-Curie grant agreement No [801215]. A. Tenesa acknowledged funding from the BBSRC through programme grants BBS/E/D/10002070 and BBS/E/D/30002275, MRC research grant MR/P015514/1, and HDR-UK award HDR-9004. O. Canela-Xandri was supported by MR/R025851/1. R.X. was supported by Australian Research Council’s Discovery Projects (DP200100499). P.N. was supported by MRC funding (MC_UU_00007/10). L.M. was supported in part by AFRI grant numbers 2020-67015-31398 and 2021-67015-33409 from the USDA National Institute of Food and Agriculture (NIFA). G.E.L., B.D.R., and C.P.V.T. were supported by appropriated project 8042-31000-001-00-D, “Enhancing Genetic Merit of Ruminants Through Improved Genome Assembly, Annotation, and Selection” of the Agricultural Research Service (ARS) of the United States Department of Agriculture (USDA). C.-J.L. was supported by appropriated project 8042-31310-078-00-D, “Improving Feed Efficiency and Environmental Sustainability of Dairy Cattle through Genomics and Novel Technologies” of ARS-USDA. J.B.C. was supported by appropriated project 8042-31000-002-00-D, “Improving Dairy Animals by Increasing Accuracy of Genomic Prediction, Evaluating New Traits, and Redefining Selection Goals” of ARS-USDA. This research used resources provided by the SCINet project of the USDA ARS project number 0500-00093-001-00-D. Mention of trade names or commercial products in this article is solely for the purpose of providing specific information and does not imply recommendation or endorsement by the USDA. The USDA is an equal opportunity provider and employer. The authors have not stated any conflicts of interest.

We thank US dairy producers for providing phenotypic, genomic, and pedigree data through the Council on Dairy Cattle Breeding under ARS-USDA Material Transfer Research Agreement 58-8042-8-007. Access to 1000 Bull Genomes Project data was provided under ARS-USDA Data Transfer Agreement 15443. International genetic evaluations were calculated by the International Bull Evaluation Service (Interbull; Uppsala, Sweden).

## Author contributions

L.F., A.T. and G.E.L. conceived and designed the project. S.L., Y.G., O.C.-X., S.W., L.F., R.X. W.C., B.J., C.X., Y.Yao, Z.Y. and X.L., performed bioinformatics analyses. O.C.-X., L.F., E.P.-C., K. D., K.R., C.-J.L., A.J.C., P. N., D.R., B.D.R., C.P.V.T., P.M.V., J.B.C., Y.Yu, S.Z. and A.T. contributed to the resource generation. S.L., Y.G., L.F., G.E.L. and A.T. wrote the manuscript. All authors read, edited and approved the final manuscript.

## Data availability statement

All raw sequencing data analyzed in this study are publicly available in NCBI Gene Expression Omnibus (GEO; https://www.ncbi.nlm.nih.gov/geo/). Details of these data can be found in Table S1-2. All processed data, the full summary statistics of QTL mapping and computational scripts can be available at https://cgtex.roslin.ed.ac.uk/.

## References

1. Buniello, A. et al. The NHGRI-EBI GWAS Catalog of published genome-wide association studies, targeted arrays and summary statistics 2019. Nucleic Acids Research 47, D1005–D1012 (2019).

2. Hu, Z.L., Park, C.A. & Reecy, J.M. Building a livestock genetic and genomic information knowledgebase through integrative developments of Animal QTLdb and CorrDB. Nucleic Acids Research 47, D701–D710 (2019).

3. Consortium, G.T. The GTEx Consortium atlas of genetic regulatory effects across human tissues. Science 369, 1318–1330 (2020).

4. Fang, L. et al. Comprehensive analyses of 723 transcriptomes enhance genetic and biological interpretations for complex traits in cattle. Genome Res 30, 790–801 (2020).

5. Xiang, R. et al. Quantifying the contribution of sequence variants with regulatory and evolutionary significance to 34 bovine complex traits. Proc Natl Acad Sci U S A 116, 19398–19408 (2019).

6. Prowse-Wilkins, C.P. et al. Putative Causal Variants Are Enriched in Annotated Functional Regions From Six Bovine Tissues. Front Genet 12, 664379 (2021).

7. Xiang, R. et al. Genome variants associated with RNA splicing variations in bovine are extensively shared between tissues. BMC Genomics 19, 521 (2018).

8. Cesar, A.S.M. et al. Identification of putative regulatory regions and transcription factors associated with intramuscular fat content traits. BMC Genomics 19, 499 (2018).

9. Littlejohn, M.D. et al. Sequence-based Association Analysis Reveals an MGST1 eQTL with Pleiotropic Effects on Bovine Milk Composition. Sci Rep 6, 25376 (2016).

10. Hayes, B.J. & Daetwyler, H.D. 1000 Bull Genomes Project to Map Simple and Complex Genetic Traits in Cattle: Applications and Outcomes. Annu Rev Anim Biosci 7, 89–102 (2019).

11. Deelen, P. et al. Calling genotypes from public RNA-sequencing data enables identification of genetic variants that affect gene-expression levels. Genome Med 7, 30 (2015).

12. Hombach, S. & Kretz, M. Non-coding RNAs: Classification, Biology and Functioning. Adv Exp Med Biol 937, 3–17 (2016).

13. Consortium, G.T. Human genomics. The Genotype-Tissue Expression (GTEx) pilot analysis: multitissue gene regulation in humans. Science 348, 648–60 (2015).

14. Tenesa, A. et al. Estimation of linkage disequilibrium in a sample of the United Kingdom dairy cattle population using unphased genotypes. J Anim Sci 81, 617–23 (2003).

15. Fang, L. et al. Functional annotation of the cattle genome through systematic discovery and characterization of chromatin states and butyrate-induced variations. BMC Biol 17, 68 (2019).

16. Dixon, J.R. et al. Topological domains in mammalian genomes identified by analysis of chromatin interactions. Nature 485, 376–80 (2012).

17. Jiang, J. et al. Functional annotation and Bayesian fine-mapping reveals candidate genes for important agronomic traits in Holstein bulls. Commun Biol 2, 212 (2019).

18. Freebern, E. et al. GWAS and fine-mapping of livability and six disease traits in Holstein cattle. BMC Genomics 21, 41 (2020).

19. Fang, L. et al. Genetic and epigenetic architecture of paternal origin contribute to gestation length in cattle. Commun Biol 2, 100 (2019).

20. Gao, Y. et al. Single-cell transcriptomic analyses of dairy cattle ruminal epithelial cells during weaning. Genomics 113, 2045–2055 (2021).

21. Kim-Hellmuth, S. et al. Cell type-specific genetic regulation of gene expression across human tissues. Science 369(2020).

22. Clark, E.L. et al. From FAANG to fork: application of highly annotated genomes to improve farmed animal production. Genome Biology 21(2020).

23. Xiang, R.D. et al. Genome-wide fine-mapping identifies pleiotropic and functional variants that predict many traits across global cattle populations. Nature Communications 12(2021).

24. Bolger, A.M., Lohse, M. & Usadel, B. Trimmomatic: a flexible trimmer for Illumina sequence data. Bioinformatics 30, 2114–20 (2014).

25. Rosen, B.D. et al. De novo assembly of the cattle reference genome with single-molecule sequencing. Gigascience 9(2020).

26. Pertea, M., Kim, D., Pertea, G.M., Leek, J.T. & Salzberg, S.L. Transcript-level expression analysis of RNA-seq experiments with HISAT, StringTie and Ballgown. Nat Protoc 11, 1650–67 (2016).

27. Liao, Y., Smyth, G.K. & Shi, W. featureCounts: an efficient general purpose program for assigning sequence reads to genomic features. Bioinformatics 30, 923–30 (2014).

28. Li, Y.I. et al. Annotation-free quantification of RNA splicing using LeafCutter. Nat Genet 50, 151–158 (2018).

29. Van der Auwera, G.A. et al. From FastQ data to high confidence variant calls: the Genome Analysis Toolkit best practices pipeline. Curr Protoc Bioinformatics 43, 11 10 1–11 10 33 (2013).

30. Browning, B.L., Zhou, Y. & Browning, S.R. A One-Penny Imputed Genome from Next-Generation Reference Panels. Am J Hum Genet 103, 338–348 (2018).

31. Chang, C.C. et al. Second-generation PLINK: rising to the challenge of larger and richer datasets. Gigascience 4, 7 (2015).

32. Patterson, N., Price, A.L. & Reich, D. Population structure and eigenanalysis. PLoS Genet 2, e190 (2006).

33. Castel, S.E., Mohammadi, P., Chung, W.K., Shen, Y. & Lappalainen, T. Rare variant phasing and haplotypic expression from RNA sequencing with phASER. Nat Commun 7, 12817 (2016).

34. Krueger, F. & Andrews, S.R. Bismark: a flexible aligner and methylation caller for Bisulfite-Seq applications. Bioinformatics 27, 1571–2 (2011).

35. Li, H. & Durbin, R. Fast and accurate short read alignment with Burrows-Wheeler transform. Bioinformatics 25, 1754–1760 (2009).

36. Ramirez, F. et al. High-resolution TADs reveal DNA sequences underlying genome organization in flies. Nat Commun 9, 189 (2018).

37. Servant, N. et al. HiC-Pro: an optimized and flexible pipeline for Hi-C data processing. Genome Biol 16, 259 (2015).

38. Song, Q. et al. A reference methylome database and analysis pipeline to facilitate integrative and comparative epigenomics. PLoS One 8, e81148 (2013).

39. Liu, H. et al. Systematic identification and annotation of human methylation marks based on bisulfite sequencing methylomes reveals distinct roles of cell type-specific hypomethylation in the regulation of cell identity genes. Nucleic Acids Res 44, 75–94 (2016).

40. Stegle, O., Parts, L., Piipari, M., Winn, J. & Durbin, R. Using probabilistic estimation of expression residuals (PEER) to obtain increased power and interpretability of gene expression analyses. Nat Protoc 7, 500–7 (2012).

41. Ongen, H., Buil, A., Brown, A.A., Dermitzakis, E.T. & Delaneau, O. Fast and efficient QTL mapper for thousands of molecular phenotypes. Bioinformatics 32, 1479–85 (2016).

42. Mohammadi, P., Castel, S.E., Brown, A.A. & Lappalainen, T. Quantifying the regulatory effect size of cis-acting genetic variation using allelic fold change. Genome Res 27, 1872–1884 (2017).

43. Wen, X., Lee, Y., Luca, F. & Pique-Regi, R. Efficient Integrative Multi-SNP Association Analysis via Deterministic Approximation of Posteriors. Am J Hum Genet 98, 1114–1129 (2016).

44. Willer, C.J., Li, Y. & Abecasis, G.R. METAL: fast and efficient meta-analysis of genomewide association scans. Bioinformatics 26, 2190–1 (2010).

45. Urbut, S.M., Wang, G., Carbonetto, P. & Stephens, M. Flexible statistical methods for estimating and testing effects in genomic studies with multiple conditions. Nat Genet 51, 187–195 (2019).

46. Shabalin, A.A. Matrix eQTL: ultra fast eQTL analysis via large matrix operations. Bioinformatics 28, 1353–8 (2012).

47. Yang, J., Zaitlen, N.A., Goddard, M.E., Visscher, P.M. & Price, A.L. Advantages and pitfalls in the application of mixed-model association methods. Nat Genet 46, 100–6 (2014).

48. Barbeira, A.N. et al. Exploring the phenotypic consequences of tissue specific gene expression variation inferred from GWAS summary statistics. Nat Commun 9, 1825 (2018).

49. Barbeira, A.N. et al. Integrating predicted transcriptome from multiple tissues improves association detection. PLoS Genet 15, e1007889 (2019).

50. Wen, X., Pique-Regi, R. & Luca, F. Integrating molecular QTL data into genome-wide genetic association analysis: Probabilistic assessment of enrichment and colocalization. PLoS Genet 13, e1006646 (2017).

51. Heger, A., Webber, C., Goodson, M., Ponting, C.P. & Lunter, G. GAT: a simulation framework for testing the association of genomic intervals. Bioinformatics 29, 2046–8 (2013).

52. Yu, G., Wang, L.G., Han, Y. & He, Q.Y. clusterProfiler: an R package for comparing biological themes among gene clusters. OMICS 16, 284–7 (2012).

53. Vavrek, M. Fossil: Palaeoecological and Palaeogeographical Analysis Tools. Palaeontologia Electronica 14, 16 (2011).

